# Holocene selection for variants associated with cognitive ability: Comparing ancient and modern genomes

**DOI:** 10.1101/109678

**Authors:** Michael A. Woodley Menie, Shameem Younuskunju, Bipin Balan, Davide Piffer

## Abstract

Human populations living in Eurasia during the Holocene experienced considerable microevolutionary change. It has been predicted that the transition of Holocene populations into agrarianism and urbanization brought about culture-gene coevolution that favoured via directional selection genetic variants associated with higher general cognitive ability (GCA). To examine whether GCA might have risen during the Holocene, we compare a sample of 99 ancient Eurasian genomes (ranging from 4.56 to 1.21 kyr BP) with a sample of 503 modern European genomes, using three different cognitive polygenic scores. Significant differences favouring the modern genomes were found for all three polygenic scores (Odds Ratios=0.92, *p*=0.037; 0.81, *p*=0.001 and 0.81, *p*=0.02). Furthermore, a significant increase in positive allele count over 3.25 kyr was found using a subsample of 66 ancient genomes (*r*=0.217, *p_one-taiied_=0.04).* These observations are consistent with the expectation that GCA rose during the Holocene.

## Introduction

The Holocene (11.7 kyr BP to the present) was a time of considerable microevolutionary change, especially among the populations of Europe and Asia. Based on the degree to which new haplotypes were arising and diverging, it has been estimated that the rate of adaptive evolution among these populations may have been 100 times greater than during the preceding Pleistocene (1). Novel adaptations that are known to have arisen and spread during this period include lactase persistence (2) and alterations in haemoglobin permitting enhanced tolerance to diminished oxygen levels among populations living at altitude (3).

Other significant adaptations that may have been favoured by selection during this period are the general learning and problem solving mechanisms that give rise to General Cognitive Ability (GCA). At the level of individual differences in cognitive performance, GCA is associated with the positive manifold that exists among performance variances on different measures of cognitive abilities. In addition to GCA, cognitive ability measures also tap specialized sources of ability variance, which are associated with performance in specific domains (e.g. verbal, spatial and perceptual abilities) (4). GCA is highly heritable with behaviour genetic twin studies indicating that between 22% and 88% of the variance in GCA, depending on age, may be due to the action of additive genetic effects (5). Consistent with this, studies employing genome-wide complex trait analysis (GCTA) GREML have found that a significant amount of the variance in GCA can be directly attributed to large numbers of variants with individually small, but collectively large (additive) effects on the phenotype (6).

GCA is not just present in human populations, but is present (at lower levels) also in a variety of animal taxa, including chimpanzees and other primates, racoons, mice, rats and ravens (7). The existence of these species differences furthermore indicates that GCA has been under directional selection, particularly in the primate clade. This seems to be especially true in the case of abilities associated with tool-use, which in primates are among the most strongly associated with GCA, in addition to being the most heritable and are also associated with the strongest signals of recent directional selection (8, 9). This is consistent with the theoretical expectation (10) that GCA is an adaptation to coping with *evolutionarily novel* challenges - these being challenges that occur infrequently throughout the phylogeny of a lineage, thus cannot be solved with recourse to specialized cognitive systems (such as those associated with cheater detection or language acquisition). Instead such problems require generalized and open-ended problem solving systems, such as learning and working memory, in order to tailor solutions to them. The ability to *innovate* a solution to such a problem (via the development of a tool) is a key manifestation of the action of these generalized problem-solving systems (10).

The Holocene is believed to have been an *environment of evolutionary relevance* for GCA as it was characterized by major transitions among human populations, especially in Eurasia, away from a hunter-gatherer mode of subsistence, towards a sedentary agriculture-based one and beyond that to urbanization (1, 11). This bought with it many evolutionarily novel problems, such as having to cope with increased population densities (1, 11). Innovations that played a major role in facilitating this transition would have included the domestication of cultivars and animals and the development of novel tools for raising the productivity of land (such as the plough) (11). Cultural innovations such as monotheism, monarchy, aristocracy, feudalism and currency-based economics arose in response to the need for coping with the hierarchical power distribution characteristic of large, static populations (11). Those populations that were successful in using innovations to solve novel problems would furthermore have had an advantage in warfare, being better able to innovate weapons and tactics, allowing them to replace less successful populations. Population growth would have increased the chances of rare GCA enhancing mutations arising, which would have favoured the aggregate fitness of those populations permitting them to expand to the greatest extent (1, 11). Thus positive *culture-gene co-evolutionary feedback* occurring during the Holocene might have rapidly increased GCA (11).

As was mentioned previously, GCTA GREML has demonstrated that GCA arises from the action of many variants each with small, but additively large effects on the phenotype (6). High-resolution genome-wide association studies (GWAS) have identified a number single nucleotide polymorphisms (SNPs), primarily associated with the development of the central nervous system, which predict variance in both GCA and educational attainment – with which GCA shares approximately 60% of its linkage pruned genetic variance (12, 13, 14, 15, 16). These SNPs can be concatenated into *cognitive polygenic scores* (henceforth POLY_COG_), which are normally distributed and can be used as a molecular index of GCA and educational attainment in standard regression-type analyses. In the present analysis, a sample of 99 ancient Holocene genomes sourced from Europe and Central Asia, aged between 4.56 and 1.21 kyr (mean age=3.44 kyr, SD=0.62 kyr), with reasonable (approximately 30%) coverage depth (17) will be used to test the hypothesis of gene frequency change, favouring higher POLY_COG_ - and therefore higher GCA. This will be achieved by comparing the aggregate POLY_COG_ levels of these ancient genomes with a genetically matched modern population, specifically the 503 individuals comprising the 1000 Genomes Phase3 EUR sample (for full details on these samples see the *Materials and Methods* section). Such an analysis builds on previous comparative analyses utilizing these same samples for tracking the changes in single SNPs associated with specific phenotypic traits (such as skin reflectance, eye colour and lactase persistence) (17).

## Results

The data were categorical (positive vs. negative allele/ancient vs. modern population) hence a 2×2 contingency table was created for each of the three POLY_COG_ (for details of how each polygenic score was computed, see the *Materials and Methods* section) assigning allele counts by GWAS effect status (positive or negative effect) to ancient and modern genomes. Odds ratios (OR) of the difference in POLY_COG_ between the ancient and modern genomes were computed and are reported in Table 1. As Fisher’s Exact Tests and *χ^2^* Tests showed nearly identical results, only the former are reported. In addition the *G*-Test (log-likelihood ratio) is utilized as an alternative test of the significance of the difference. As a further robustness test, the difference in POLY_COG_ between the two sets of genomes was examined using the unconditional Barnard’s and Boschloo’s Exact Tests. These statistics are extremely computationally intensive when large numbers of pair-wise comparisons are employed. Therefore they could only be used for the two smaller POLY_COG_ (9 and 11 SNP). The results were identical to those obtained via the conditional techniques.

**Table 1.** 2 × 2 contingency tables with positive and negative GWAS effect allele counts for ancient and modern genomes for each of the three POLY_COG_, along with Fisher’s Exact Test OR and *G*-Test. The non-integer allele counts result from averaging the genomes of two samples (RISE507 and RISE508) that came from the same individual. The samples had different allele counts likely resulting from different coverage.

| **POLY<sub>COG</sub>** | **Genome grouping** | **Positive allele count** | **Negative allele count** | **Fisher's Exact Test. OR; 95%CI; *p*-value.** | ***G*-Test; *p*-value** |
| --- | --- | --- | --- | --- | --- |
| **Nine linked-hits** | Ancient genome | 177 | 273 | 0.92 (0.665-0.990); | *G*=4.445; |
| **POLY<sub>COG</sub>. *N*=9** | Modern genome | 4017 | 5037 | *p*= 0.037 | *p*=0.035 |
| **Pooled analysis** | Ancient genome | 3298.5 | 3997.5 | 0.81 (0.882-0.969); | *G*=10.486; |
| **POLY<sub>COG</sub> (Okbay et al., 2016). *N*=130** | Modern genome | 61666 | 69114 | *p*=0.001 | *p*=0.001 |
| **Significant hits** | Ancient genome | 268.5 | 282.5 | 0.81 (0.683-0.969); | *G*=5.536; |
| **POLY<sub>COG</sub> (Davies et al., 2016). *N*=11** |  |  |  | *p*= 0.02 | *p*=0.019 |

A forest plot of the OR values for the three POLY_COG_ scores is presented in Figure 1.

**Fig. 1.**
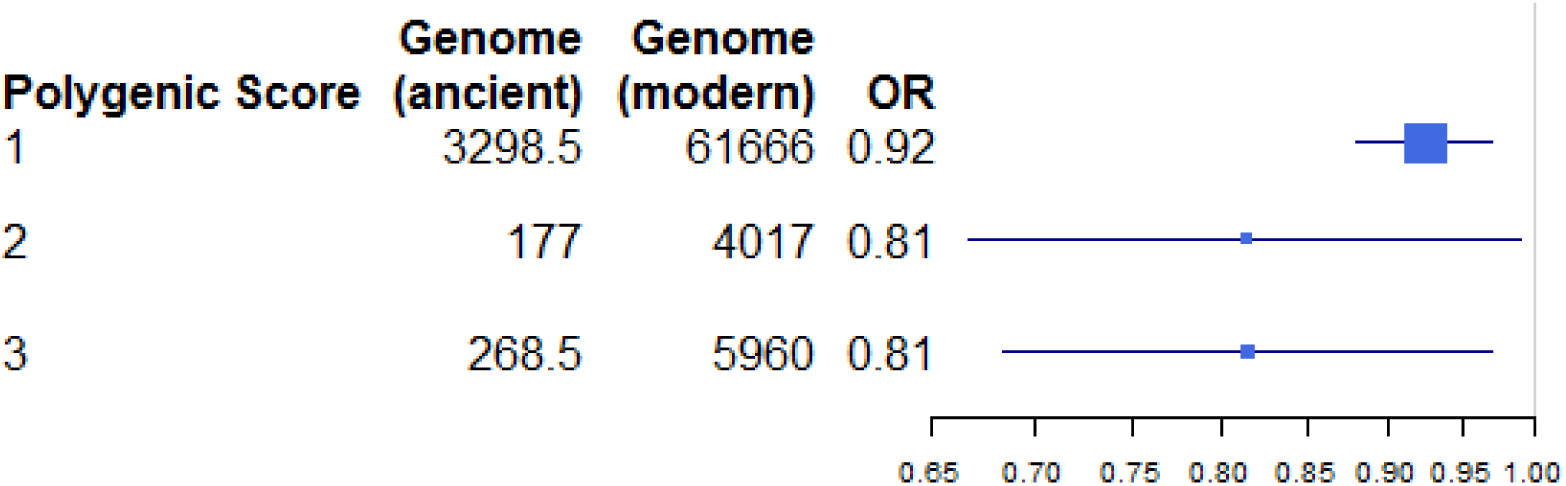
Forest plot of the OR of the difference between ancient and modern genomes for three POLY_COG_ (130, 9 and 11 SNP, labelled 1, 2 and 3 respectively). Box size corresponds to SNP numbers.

These results indicate that modern European genomes have higher POLY_COG_ relative to ancient ones (sourced from Europe and Central Asia).

A second analysis was conducted to investigate the association between POLY_COG_ and sample year within the subset of 66 ancient genomes for which age estimates were available (17). If selection is operating on these variants throughout the 3.25 kyr covered by this sample then less ancient genomes should have higher POLY_COG_ relative to more ancient ones. The correlation of positive allele frequency with the sample year (scaled using the BCE/CE calendar era) was in the expected positive direction (samples from more recent years had higher positive allele counts) the result is significant when a one-tailed test is used (*r*=0.217, 95% CI=-0.026 to 0.043, *p_one-tailed_*=0.04), which is theoretically justified on the basis that the direction of the effect was anticipated (18). The scatter plot is presented in Figure 2.

**Fig. 2.**
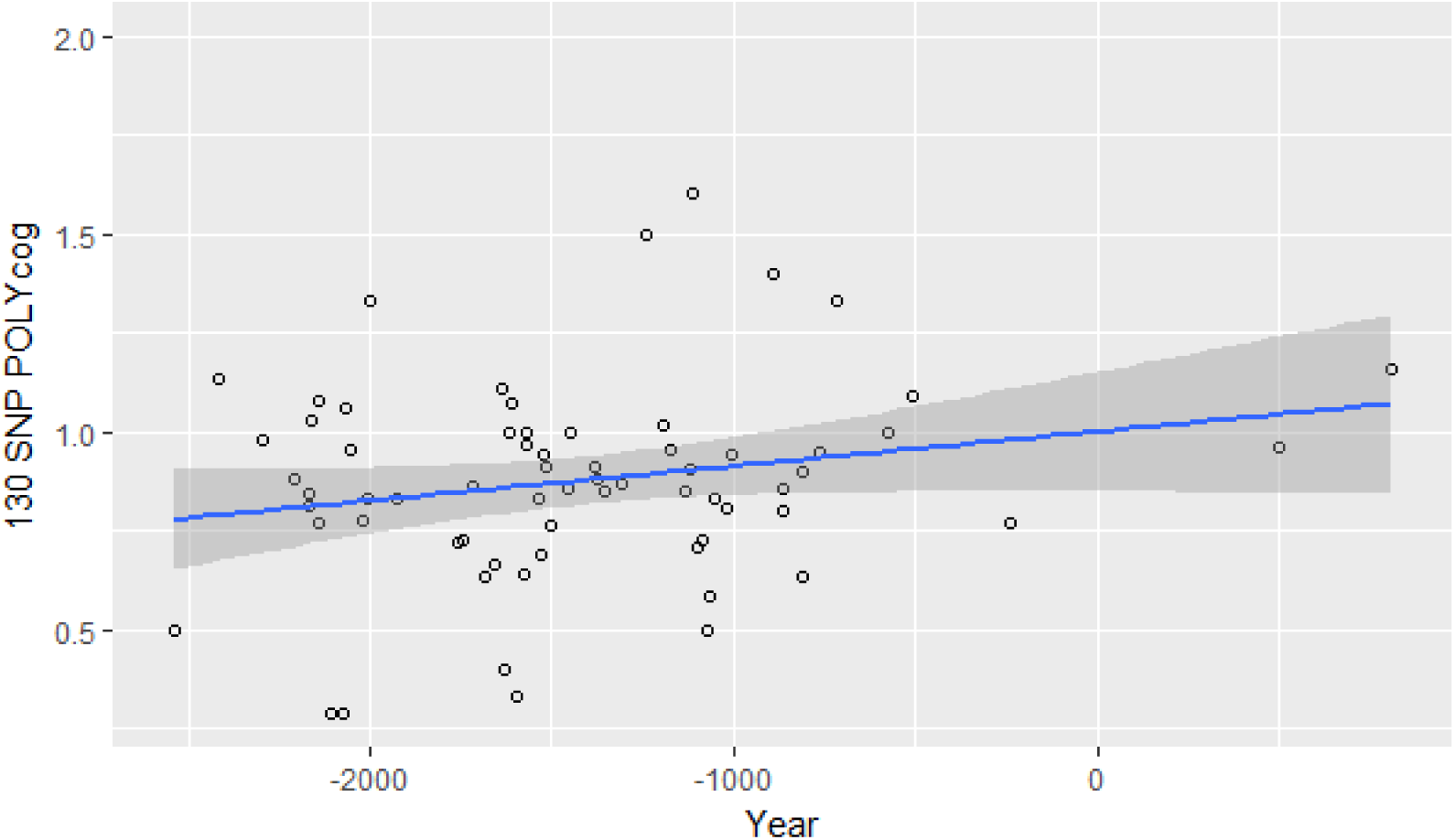
Scatter plot of 130 SNP POLY_COG_ positive allele counts for 66 ancient genomes as a function of year (scaled in terms of the BCE/CE calendar eras). *r*=0.217, *p_one-tailed_=0.04, N=66.* Grey area around trend line corresponds to the 95% CI.

## Discussion

Consistent with theoretical expectations, the Holocene appears to have been an environment of evolutionary relevance for GCA, as modern European genomes have higher POLY_COG_, relative to those sourced (predominantly) from Bronze Age Europe and Central Asia. A second analysis revealed that within a subset of the ancient genome sample, there was a (one-tailed) significant trend towards increased positive allele counts over time.

This increase in POLY_COG_ can be theoretically accounted for in four different ways: i) the appearance of new mutations, ii) genetic drift involving already standing genetic variation, iii) microevolutionary selection pressures on standing genetic variation, resulting in soft sweeps (19, 20), and iv) population expansion, replacement and admixture due to migration.

The first two theories can be discounted, as the POLY_COG_ scores utilized here are comprised of SNPs that are present (either directly or in strong linkage) in both the ancient and modern populations. Therefore these particular SNPs did not arise *de novo* in this period. Genetic drift based models would make the assumption that the variants comprising POLY_COG_ are selectively neutral, and that their distributions should therefore correspond to the action of genetic diffusion and population bottlenecks. It is unclear however why this would lead to modern genomes being higher in POLY_COG_ relative to ancient ones, and is furthermore at odds with the observation from studies utilizing various alternative POLY_COG_ that these do not appear to be selectively neutral in contemporary human populations (21, 22, 23, 23, 25, 26). This leaves selection pressure acting on standing genetic variation and population expansion, replacement and admixture.

The idea that selection takes many millennia to produce noticeable changes in a population is at odds with data indicating that within human populations, changes in the means of traits such as height and age at menarche can occur due to selection in as little as five decades (27). Selection operating over the course of millennia (as in the present case) would be expected to produce quite considerable microevolutionary change. As was discussed in the *Introduction,* Holocene populations, especially those in Eurasia, appear to have undergone accelerated adaptive microevolution relative to those living in the Pleistocene (1, 11). Increasing cultural complexity and technological sophistication among Holocene populations therefore likely arose in part from selection favouring GCA. Cultural and technological change can in turn create, via feedback, conditions favouring higher GCA (11, 28). This process likely continued until the Late Modern Era, where it has been noted that among European populations living between the 15^th^ and early 19^th^ centuries, those with higher social status (which relates to GCA via common causal variants; 29) typically produced the most surviving offspring. These in turn tended towards downward social mobility due to intense competition, replacing the reproductively unsuccessful low-status stratum and effectively ‘bootstrapping’ those populations via the application of high levels of skill to solving problems associated with production and industry, eventually leading to the Industrial Revolution in Europe (30, 31). The millennia long microevolutionary trend favouring higher GCA not only ceased, but went into reverse among European-derived populations living in the 19^th^ century (32), largely in response to factors such as the asymmetric use of birth control and prolonged exposure to education among those with high GCA (33). Consistent with this, it has been found that various POLY_COG_ negatively predict fertility in contemporary Western populations (23, 24, 25, 26). It is important to note that this microevolutionary process (working in the opposite direction) has likely attenuated the difference in POLY_COG_ between the modern and ancient genomes noted in the present study.

Changes in allele frequencies can also occur via population expansion and replacement. The end of the Last Glacial Maximum initiated major cultural changes, leading to the development of agriculture and the subsequent Neolithicization process (8-5 kyr BP), which involved transmission both of cultural and genetic factors spreading from the Middle East (34). Another big cultural change associated with major population movements was the Bronze Age, whose culture started replacing the Neolithic farming cultures in temperate Eastern Europe 5 kyr BP (17). This *Pontic-Caspian Steppe* genetic component among contemporary Europeans is associated with the Corded Ware and Yamnaya people, and possibly accounts for the spread of Indo-European languages (17). Being potentially highly fitness enhancing, admixture among populations may have furthermore led to the constituent variants of POLY_COG_, moving rapidly between populations. However recent evidence suggests that the European and Central Asian gene pools towards the end of the Bronze Age closely mirror present-day Eurasian genetic structure (17). Hence, population movements and replacement likely played a larger role in the evolution of GMA mostly prior the period covered by the present sample. Sicily and Sardinia, and the southern fringes of Europe in general are notable exceptions however, as they tend to cluster genetically with Neolithic farmers (17).

The third theoretical scenario (microevolutionary change) is therefore the most likely explanation for the POLY_COG_ increase during the time period covered by our sample.

It has been noted that concomitant in time with the apparent increase in POLY_COG_ observed presently is a parallel selection trend favouring smaller brains (35). These trends may at first appear to be contradictory, as GMA is positively associated with brain volume. It must be noted however that meta-analysis reveals that the association is weak (*ρ*=0.24), indicating that the majority of the variance in brain volume is unrelated to variance in GMA (36). Furthermore studies utilizing POLY_COG_ have found that it does not predict variation in brain volume, despite both POLY_COG_ and brain volume making independent contributions to GMA (37). These findings suggest that the decline in brain volume during the Holocene may have been a consequence of enhanced brain efficiency stemming from increased corticalization and neuronal connectivity, with more bioenergetically optimized brains simply requiring less mass to achieve greater processing power. It may therefore be this process that the increase in POLY_COG_ is tracking in our samples. A prediction stemming from (35) is that variants that predict brain volume and not GMA should show the opposite trend in time to POLY_COG_ when examined in the context of the present samples.

A limitation on the present work that needs to be discussed is linkage disequilibrium (LD) decay. This takes place when a pair of SNPs on a chromosome in a population move from linkage disequilibrium to linkage equilibrium over time, due to recombination events eventually occurring between every possible point on the chromosome (38). LD decay can be an issue when comparing polygenic scores computed using genetically different populations (i.e. either across time or space). Since most GWAS hits are actually tag SNPs, decay in LD implies that the causal SNPs will be further removed from the tag SNPs. However, the tag SNPs will resemble a sample of random SNPs. Since the frequency of the average SNP allele is 50%, the GWAS hits will tend to converge towards an average frequency of 50%, with increasing LD decay. The implication of this for our analysis is that our estimates of the difference in POLY_COG_ are therefore conservative, when these have a frequency lower than 50%, whilst the opposite is true when they have higher frequencies (the average frequencies in the modern 1000 Genomes samples were the following: for the 130 SNP POLY_COG_ = 47%, 9 SNP POLY_COG_ = 44% and the 11 SNP POLY_COG_ = 53%). Furthermore, because GCA likely involves around 10,000 causal variants (39), and as there is relatively little genetic distance between the ancient and modern genomes, as indicated by the *F_st_* value (0.016), the results of simulations indicate that LD decay is ultimately expected to be minimal (40) and should not be confounding the results.

The present effort constitutes a purely empirical attempt to show the theoretically anticipated change in POLY_COG_ over time utilizing very simple methodology, which makes few modelling assumptions. The difference in POLY_COG_ between the ancient and modern genomes is indifferent to the use of alternate versions of POLY_COG_ and to the use of various statistical techniques for quantifying the significance of the group difference. This suggests that the finding enjoys multi-trait, multi-method validity (41). Building on this, future research could examine the spread of POLY_COG_ via admixture analysis using ancient genomes. Currently this has been used to track single SNPs (17), tracking complex polygenic scores is doubtlessly more of a challenge however, requiring bigger samples and also analytical techniques that are sensitive to changes in the linkage phase of the SNPs comprising the polygenic scores over time.

## Materials and Methods

### Modern genomes

Previous studies comparing ancient and modern genomes have utilized the Phase3 1000 Genomes dataset (42) as a reference set, on the basis that it is the most globally representative set of genomes currently available (17). The 2,504 samples in the Phase3 release are sourced from 26 populations, which can be categorised into five super-populations by continent. These are East Asian (EAS), South Asian (SAS), African (AFR), European (EUR) and American (AMR). These genomes were obtained in the form of VCF files, via the online 1000genomes FTP. The EUR sample, which is the most closely related to the ancient genomes (17), is comprised of 503 individuals from five different countries (Finland, Great Britain, Italy, Spain and the United States – specifically North-Central Europeans from Utah).

### Ancient genomes

Data on the ancient Holocene genomes were obtained from the *European Nucleotide Archive* in the form of BAM files (accession number PRJEB9021). These genomes, which were used previously to analyse the spread of variants associated with skin and eye colour, among other traits (17), were aged between 4.56 and 1.21 kyr (mean age=3.44 kyr, SD=0.62 kyr), covering the early Bronze to early Iron ages. There are 102 genomes in total, however two samples (RISE507 and RISE508) were sourced from the same individual (17). The total allele count was therefore calculated using the full sample alternately using each version of the genotype then the results of the two analyses were averaged. The majority of the ancient genomes were sourced from regions that are presently considered to be parts of Europe, the remainder being sourced from Central Asia. Late Bronze Age European and Central Asian gene pools resemble present-day Eurasian genetic structure (17). Indeed, with values of *F_st_* ranging from 0.00 to 0.08, there is little to modest genetic differentiation between the present-day 1000 Genomes EUR and the Ancient samples (little differentiation corresponds to an *F_st_* range of 0 to 0.05, and modest to an *F_st_* range of 0.05 to 0.15 [43]). These values are lower than the distance between present-day Europeans and East Asians (*F_st_*=0.11) (17). Despite this the two ancient genomes belonging to the Siberian *Okunevo* culture (RISE515 and RISE516) were somewhat of an outlier, exhibiting modest differentiation relative to the EUR sample when compared with the other genomes in the sample (average *F_st_*=0.074 vs. 0.016 for the remainder of the sample). Their removal therefore reduced the genetic differentiation between the two samples, yielding 99 ancient genomes, sourced from sites located in present-day Armenia (8.08%), Czech Republic (6.06%), Denmark (6.06%), Estonia (1.01%), Germany (10.1%), Hungary (10.1%), Italy (3.03%), Kazakhstan (1.01%), Lithuania (1.01%), Montenegro (2.02%), Poland (7.07%), Russia (36.36%) and Sweden (8.08%).

### Reference genome

Paired-end reads are aligned to the reference human genome obtained (in the form of a FASTA file) from the UCSC database (GRCh37/hg19).

### Read realignment and base recalibration

After removing duplicate reads the reads were re-aligned around the known indels from the 1000-genome sample using the *GenomeAnalysisTKLite-2.3-9* toolkit. The known indels set was obtained from the GATK resource page (44). After performing realignment the base re-calibration step is performed, as is recommended (45, 46). After recalibration, the quality score of each base becomes more accurate. Known variant position is taken into account to recalibrate the quality score.

### Variant calling and computing POLY_COG_

After performing re-alignment the GenomeAnalysisTKLite-2.3-9 toolkit *UnifiedGenotyper* was used to identify the single nucleotide variants (SNPs) and short indels. The positive allele counts and allele frequencies of the target SNPs were calculated from the Ancient genome VCF files and also from the 1000 Genome VCF files using a custom made PERL script (available on request). Variant calling was used to construct a POLY_COG_ score from a list of nucleotides that were found to be genome wide significant ‘hits’ for educational attainment in a recent large-scale meta-analytic GWAS study, which predicted 3.6% of the variance in GCA (14). After excluding those variants that were completely absent from the ancient genomes (likely due to coverage artefacts) the resultant POLY_COG_ score consisted of 130 ‘hits’ that were present to some degree in both the ancient and modern genomes. A second POLY_COG_ score was constructed using data from a GCTA GREML study of GCA and related phenotypes (6). In this study 1,115 SNPs reached GWAS significance, of which 15 were independent signals. Four SNPs were absent from the ancient genomes thus a POLY_COG_ was calculated using 11 SNPs. A third POLY_COG_ score was constructed utilizing a technique developed in (20, 21), employing only the nine GWAS hits from (14) that were in close linkage disequilibrium (linkage cut-off *r*≥0.8, 500kb linkage window) with ‘hits’ predicting educational attainment across two other large GWAS studies (6, 12). Linkage was determined using the NIH *LDLink program* (47) with the 1000 Genomes Phase3 CEU population as a reference group. These are presented in Table 2. By focussing on only the smaller numbers of linked replicates, the third POLY_COG_ score was developed as a means of negating the likely high numbers of false positive SNPs present in larger POLY_COG_ scores. This approach is based on maximizing the signal to noise ratio of directional selection by focusing on the GWAS hits with the highest significance (or replication rate across publications), as opposed to maximizing the amount of variance explained, which instead is the aim of the classical, within population GWAS design.

**Table 2.** Nine linked replicated SNPs with linkage disequilibrium (*D’*) and *R*^2^ coefficients.

| *Publication* | *Index SNP* | *Publication SNP* | *Linked SNP source* | *D'* | *R<sup>2</sup>* |
| --- | --- | --- | --- | --- | --- |
| Davies et al., 2016 | rs12042107_C | rs1008078_C | Okbay et al., 2016 | 1 | 0.8 |
| Rietveld et al., 2013 | rs11584700_G | rs11588857_A | Okbay et al., 2016 | 1 | 0.94 |
| Rietveld et al., 2013 | rs4851266_T | rs12987662_A | Okbay et al., 2016 | 1 | 1 |
| Davies et al., 2016 | rs13086611_T | rs148734725_A | Okbay et al., 2016 | 1 | 1 |
| Davies et al., 2016 | rs11130222_A | rs11712056_T | Okbay et al., 2016 | 1 | 0.98 |
| Davies et al., 2016 | rs55686445_C | rs62263923_G | Okbay et al., 2016 | 1 | 0.98 |
| Davies et al., 2016 | rs12553324_G | rs13294439_C | Okbay et al., 2016 | 1 | 0.98 |
| Davies et al., 2016 | rs4799950_G | rs12969294_G | Okbay et al., 2016 | 1 | 0.92 |
| Rietveld et al., 2013 | rs9320913_A | rs9320913_A | Okbay et al., 2016 | 1 | 1 |

### Analysis

Statistical analysis was conducted using R v. 3.3.2 (48). In order to test our hypothesis that POLY_COG_ had different proportions between the two populations against the null hypothesis that there would be no differences, we used Fisher’s Exact Test and *χ^2^* tests, since the data are categorical (positive vs. negative allele/ancient vs. modern population). At the same time, the contingency table dealt with unequal representation of genomes for each SNP within the ancient group (i.e. some SNPs were called if they were present in only one individual) by incorporating this information (each SNP is assigned a weight proportional to the number of genomes in which the SNP was called) Fisher’s Exact Test was preferred over the *χ^2^* Test because it allows exact calculation of the deviation from the null hypothesis, rather than relying on an approximation. An alternative to Fisher’s Exact Test is the log-likelihood ratio *G*-Test, which is employed here as an additional test of significance. This was implemented using the R package *DescTools* (49). Analyses were also run for the two smaller POLY_COG_ (11 and 9 SNPs) using two unconditional tests (considered more powerful alternatives), Barnard’s (50, 51) and Boschloo’s Tests (52), utilizing the R package *Exact* (53). A second analysis was carried out on a subsample of 66 ancient genomes for which sample age was known (age range: 1208 to 4.46 kyr). The correlation was computed between individual positive allele counts derived using the 130 SNP POLY_COG_ with genome year, in order to test for change over time. Since on average the ancient genomes had about 30% coverage, carrying out this analysis using the smaller POLY_COG_ would add too much noise and produce unreliable estimates for individual genomes. Two additional genomes for which age data were present (RISE413 and RISE42) were identified as having significantly outlying values of positive allele count based on the outlier-labelling rule (54) because there was only one SNP per genome. For the analysis only genomes with at least two SNPs were retained. Removing these two outlying genomes also reduced the skew of the regression residual from 1.16 to 0.698, making the data more amenable to parametric regression.

